# An image-computable characterization of the non-conditioned linkage of visual drive and valence in the primate amygdala

**DOI:** 10.64898/2026.05.22.726311

**Authors:** Alina Peter, Gwangsu Kim, James J. DiCarlo

## Abstract

The amygdala is a key node in linking high-level representations of visual stimuli to affective state, and much research has focused on its role in learning artificial stimulus-value associations. By comparison, the primate amygdala’s encoding of natural, non-conditioned visual stimuli is less well understood. Here, we report that some amygdala neurons have visual selectivity that is systematically linked to their valence tuning, without laboratory conditioning. First, electrophysiological recordings revealed that the firing rate responses of many amygdala recording sites are selective across arbitrary non-conditioned natural visual stimuli. Second, this selectivity was well-predicted by contemporary, image-computable models of the visually-driven selectivity of high-level ventral stream neurons that provide major afferents to the amygdala. Third, a subpopulation of visually selective amygdala units also coded valence, as defined in prior work. Fourth, these valence preferences were correlated with their visual tuning, in that the image-computable models predicted which visual stimuli tended to drive positive-valence sites and which tended to drive negative-valence sites. Taken together, these results suggest that the visual drive provided from the ventral visual stream into the amygdala is strong, and that it is not unlinked or randomly linked to valence coding. Furthermore, the results establish baseline image-computable models of visual encoding in the primate amygdala.

**Significance:** Our understanding of how the primate amygdala links natural visual input to affective state remains limited. Here, we report that visual selectivity in the amygdala is not random but is systematically coupled to valence encoding, even without explicit conditioning. Amygdala neurons exhibit reliable visual selectivity for natural stimuli, and this selectivity is predicted by image-computable models of the upstream ventral visual system. Interestingly, a subpopulation of these visually selective units also coded valence, and their valence preference correlated directly with their visual preferences. This establishes a predictive model of visual and valence encoding, suggesting that visual information is wired to influence affective processing and potentially enabling noninvasive modulation of limbic circuits.

## Introduction

Humans and non-human primates can readily assign valence (positive or negative value) to visual stimuli [1]. This capacity is thought to be critical for survival, supporting both defensive and appetitive behaviors [2]. Computational studies suggest that successful visual–valence processing relies on sophisticated visual feature analysis, likely supported by the visual system [3, 4]. Yet, a neurally mechanistic account of how vision and valence are linked, which fully maps the pathways from image pixels to valence, is still largely unclear.

The amygdala is thought to be a key neural substrate for associating visual inputs to their valence [5, 6]. In primates, neurons in the basolateral amygdala receive direct projections from inferotemporal cortex (IT) [7], the culmination of the ventral visual stream (VVS), which conveys visual representations that can directly support tasks such as visual object recognition [8, 9]. Consistent with this anatomical connection, amygdala neurons show complex visual tuning with preferred responses to a broad range of images, from social and affective stimuli (e.g., faces) to neutral objects and artificial stimuli [10–12]. In parallel, so-called “valence cells” in the amygdala – those responding to positive (e.g., juice) or negative (e.g., airpuff, foot shock) events – are closely intermingled in both central and basolateral amygdala complexes [5, 13]. These neurons not only carry valence-related information, they also can play a causal role in behavior, as holographic optogenetic reactivation of cells activated during valence events is sufficient to induce appetitive versus avoidant behaviors in rodents [14].

Furthermore, seminal studies showed that amygdala neurons can encode the learned value of visual stimuli, such that neurons can respond higher (or lower) to images conditioned with a particular outcome valence (e.g. reward), and can reverse their responses when the associated outcome valence is reassigned [13, 15]. Given this prior work and related contemporary studies in rodent amygdala [5, 16, 17], a widely held view is that the amygdala is a key node in creating arbitrary, novel associations – presumably implemented as mechanistic synaptic linkages – between visual stimuli that are temporally associated with positive or negative valence outcomes.

In that context, the “adult background” state of those linkages might be thought of as having valence-neutral stimulus regimes that can be sculpted by laboratory-conditioned experience. Experimental stimuli that are as valence-neutral as possible, such as fractals, are often chosen to try to be near such regimes [13, 15]. However, evidence also suggests that some visual stimuli (e.g. snakes, faces, etc.) are already linked with valence, even without laboratory conditioning (see below). A more comprehensive understanding of these non-laboratory-imposed, “adult background” stimulus-to-valence linkages (whether learned or innate, see discussion), is needed. That goal provokes elementary questions: how are arbitrary visual stimuli encoded in amygdala neural populations? Does the adult primate amygdala carry a background (i.e. non-laboratory-conditioned) set of visual-to-valence linkages? And if so, can we characterize those linkages all the way from image pixels to their valence coding?

Based on evolutionary accounts, amygdala circuits should preferentially encode visual features that are reliable predictors of ecological value, allowing automatic value inference (i.e. valence) for novel stimuli that share these diagnostic features [1, 16, 18]. Supporting this idea, some amygdala neurons that respond more to negative unconditioned stimuli (airpuff) than positive ones (juice) also respond more to a threatening naturalistic visual cue (direct gaze of an intruder) than a nonthreatening one (averted gaze) and reward-responsiveness correlates with responsiveness to social rank[19, 20]. This suggests that some valence-tuned amygdala neurons may exhibit stable visual feature preferences not induced by laboratory conditioning. However, it is unclear whether these linkages to valence coding in the amygdala are limited to these highly specific social stimuli. More specifically, we do not yet know how to accurately predict the valence linkage of arbitrary visual stimuli.

To answer this question, we combined monkey electrophysiological recordings of neurons in the monkey amygdala with computational modeling of their responses to arbitrary visual stimuli and standard valence stimuli (water, juice and airpuff). We reasoned that, given recent advances in our field’s ability to accurately model and predict the visual responses along the ventral visual stream to arbitrary visual images, we might be able to readily extend those same models to explain and accurately predict the visually-driven responses of amygdala neurons to arbitrary visual images. Specifically, a family of deep, convolutional, task-optimized ANN models now provide image-computable accounts of neural representations at all four major stages of the ventral visual stream (V1, V2, V4, and IT), enabling quantitative prediction and model-guided, visually-driven control of neural responses at the spatial scale of single neuronal sites, including its highest area, IT [21–23]. Given the direct IT-to-amygdala axonal connections, we reasoned that amygdala visual responses may be accurately modeled as an extension of these same ventral stream models – for example, by adding one additional model layer beyond the IT model layer.

Indeed, we here report that many primate amygdala neurons exhibit reliable visual selectivity that is reasonably-well modeled by extensions of image-computable ventral stream models. Moreover, for the subset of visually-driven sites that also encode valence, we find that a site’s visual selectivity is systematically linked to its valence preference, even without explicit laboratory value conditioning.

## Results

### Reliable responses in primate amygdala to natural images

Neural activity was recorded from three monkeys performing a fixation task, with rapid serial visual presentation (RSVP) of images (Figure 1A). 400 different naturalistic images belonging to 16 different categories were presented pseudorandomly, with 7–12 images per trial, each presented for 100 ms with an inter-stimulus interval of 100 ms (median 20/27/45 repetitions (M1/M2/M3) repetitions per image across the session, see Methods). The RSVP paradigm is frequently used when studying the visually-driven responses of ventral stream neurons to efficiently assess responses to a large image set. The chosen categories reflected those believed to be of particular interest to the field (i.e. both amygdala and IT researchers [10–12, 25–27]), such as human and monkey faces, snakes, spiders, and food, and natural scenes, as well as more arbitrary categories such as chairs, tractors and beetles. In monkey 1, amygdala activity was recorded from a total of 371 single- and multi-units (215 MUA, 156 SUA), targeting the lateral aspects of the basolateral complex (Figure S1B). We will refer to these recording sites as amygdala sites rather than basolateral amygdala sites because we cannot exclude that cells from other amygdala regions are included in the sample. In monkeys 2 and 3, activity was recorded from 141 and 64 multi-unit sites respectively (Figure 1B, see Methods). For simplicity, we will refer to individual “sites” to mean both single- and multiunit activity, though we may sometimes make explicit reference to a cell or neuron, i.e. single unit (e.g. Figure 1C).

**Figure 1:**
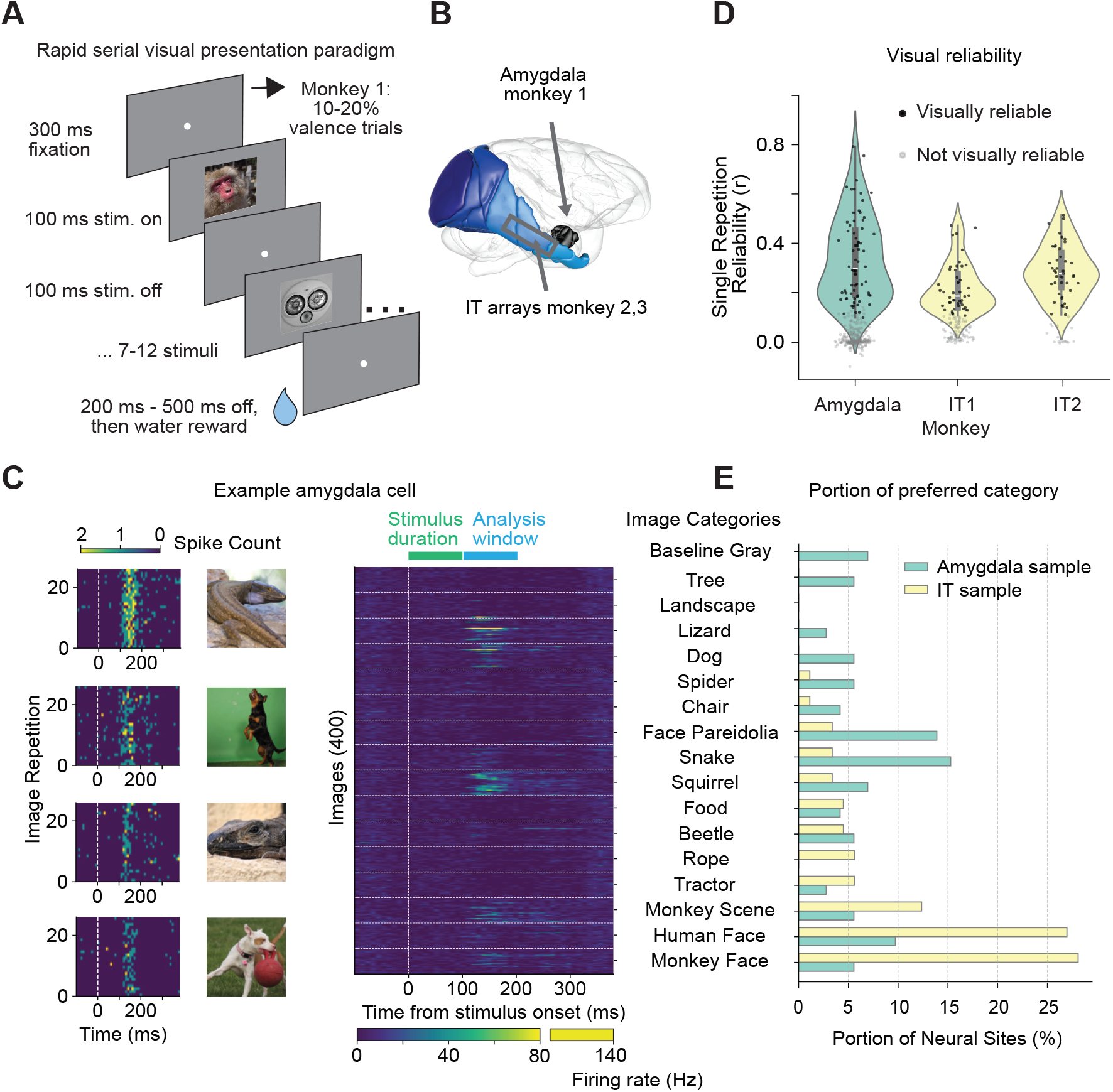
A) Task. Images were presented to fixating monkeys in randomized order, in rapid sequence, with 7–12 images per trial followed by water reward. For valence assessment in monkey M1 (amygdala recordings), in 10–20% of randomly selected trials, a tone and delay period were followed by either juice reward, water reward or airpuff. Fixation was required for all trials. See also Methods and Figure S1A. B) Illustration of recording locations. 3D representation of recording locations in amygdala (M1) and IT (M2-M3) Image generated using Scalable Brain Atlas, [24]. Figure S1B: MRI of M1. C) Left: Single-trial responses of example neuron for 4 top-driving images. Right: Per-image trial-averaged responses of example neuron. Horizontal lines indicate category boundaries. D) Reliability of visual responses across repetitions (see Methods) per animal. E) Percentage of neurons responding strongest to each category in amygdala and IT monkeys, based on cross-category average response. Figure S1D: Same based on category of strongest-driving image overall.

We first observed that some amygdala neurons exhibited reliable, short-latency responses to natural images in the RSVP paradigm. Figure 1C shows peristimulus time histograms (PSTHs) of an example single unit in the amygdala. The PSTHs of this unit to repeated presentations of the four top-driving images demonstrate consistent, image-dependent responses, with a typical latency of just over 100 ms (Figure 1C, left). Moreover, visual inspection of average responses of this unit over time for all images (Figure 1C, right) suggests stronger activity for images within certain categories (e.g., lizard and dog).

Examination of the reliability of the visually-driven selectivity of all the recorded neural sites revealed that many amygdala neurons are reliably selective, and that reliability of the recorded amygdala population is comparable to the recorded IT populations (Figure 1D). Specifically, we quantified the reliability of the visual selectivity of each site over the set of images as the mean Pearson correlation between two randomly selected repeats of responses to all 400 images, averaged over 1,000 resamplings (“Single repetition reliability”, SRR; see Methods). A unit was considered visually reliable if its SRR was significantly above zero. As shown in Figure 1D, both the amygdala and IT recordings contained a significant proportion of visually reliable units (black dots, *∼* 20% of amygdala units (72/351, 41 SUA and 31 MUA) and *∼* 43% of IT units (89/205) met this criterion). The reliability scores of visually reliable amygdala units were on par with those of visually reliable IT units, with highly overlapping distributions. Because different animals and recording techniques were used, we refrain from direct comparison via statistical analysis. Furthermore, we note that SRR in the amygdala was stable during the recording sessions, average SRR was *∼* 0.37 with random repeats and *∼* 0.36 with one repeat drawn from the first and second half of the session respectively (ca. 1 hour apart, see Methods). Grand average responses to images are shown in Figure S1C, indicating similar or somewhat longer latencies in the amygdala. In sum, we find that many amygdala sites are reliably driven by somewhat arbitrary batteries of natural visual stimuli, even during passive fixation and without any explicit value conditioning of these stimuli.

Furthermore, amygdala neurons exhibited complex visual feature preferences, reminiscent of IT neurons (see also Figure 2B for more examples, [10]). For each neuron, the preferred category was identified as the stimulus category that evoked the highest mean response (Figure 1E). Individual units in the amygdala showed preferences for a broad range of categories. In contrast, IT units showed a more pronounced, though not exclusive, preference for faces of both monkeys and humans. We also note that some amygdala units appear to prefer certain visual features (e.g., animal side views in Figure 1C), which were not reflected in the predefined object categories. Our analyses show that some neural sites in the lateral amygdala exhibit high visual reliability and intricate tuning properties.

**Figure 2:**
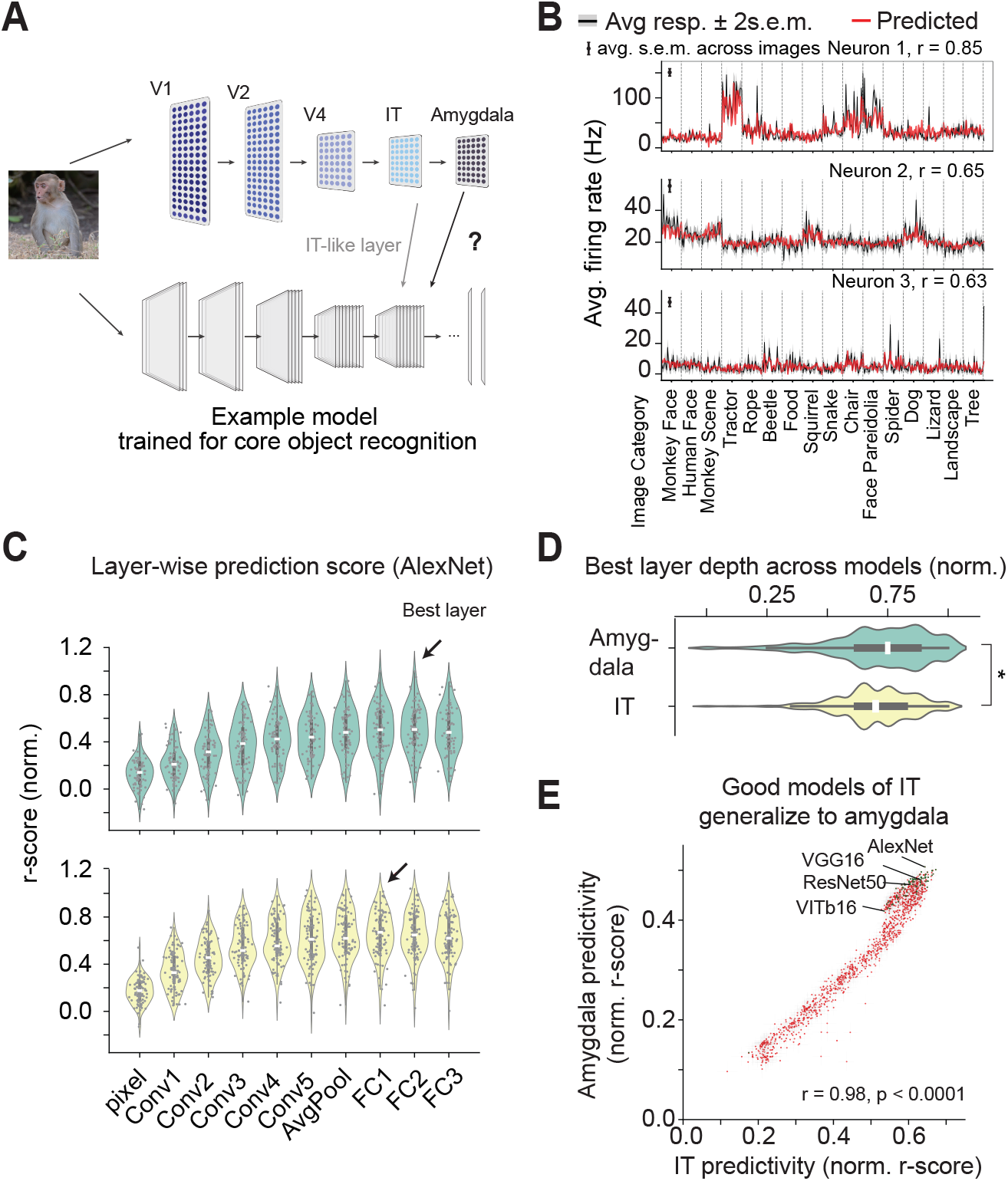
A) Illustration of the mapping of single neuron responses to layers of a deep neural network. B) Illustration of image-by-image average responses of three example cells (black lines) overlaid with the model prediction from the best-performing layer in C for IT (red lines). C) By layer fit for each visually reliable site in the amygdala (top) and IT (bottom) in an example deep neural network (AlexNet, [28] see Methods). Explained variance increases with depth. D) Peak explained variance of amygdala neural responses occurs later compared to IT, median depth amygdala 0.75, median depth IT 0.7. See Methods. E) Testing 78 different deep neural network types (see Methods, [29]) reveals that networks that show better performance in IT tend to show better performance in the amygdala, p *<* 0.0001. This relation also holds when only considering the best-performing layers per model (green dots).

### Neural models of the ventral visual stream predict amygdala neural responses

Given its direct input from IT, we hypothesized that the responses of the reliably visually-driven amygdala sites to natural images could be modeled using contemporary, machine-executable models of the VVS (Figure 2A, [8, 21]). Key parameters of these models are typically determined by optimizing the overall model performance on visual object recognition with natural images and then freezing those parameters. Models are then “mapped” onto recorded VVS neurons via linear regression methods, and prediction accuracy is tested on the responses to held-out images. Previous work has demonstrated that model neural units in early layers are good predictors of the responses of neurons in early VVS areas such as V1, while model units in deeper layers are good predictors of the responses of deeper areas of the VVS (V4 and IT) [21, 30–32]. Notably, many of the leading VVS models have one or more layers that are even deeper than the best mapped IT model layer.

To test the hypothesis that VVS models might readily extend to explain the responses of visually-driven amygdala neurons, we used the same techniques outlined above to explore a set of leading VVS models (see Methods). We were particularly interested in the model IT layers and layers beyond IT. Specifically, for any given base model and any given layer in that model, we measured how well visual responses of amygdala sites to held-out images could be predicted based on linear combinations of responses of units (linear response predictivity, fitting with cross-validated ridge regression, see Methods).

To illustrate this, we show the prediction of held-out responses to images for three example amygdala sites, of one VVS model and one model layer (the one best-performing in IT). Figure 2B shows model predictions from the FC1 layer of AlexNet [28] that has been mapped to these three sites. We note that the predicted responses (red) closely follow the measured firing rates across 400 images (black; data never used for mapping). Because this model layer is also a reasonably good predictor of IT responses (see Figure 2C), this suggests that visual representations useful for modeling IT responses can also at least partially account for amygdala responses. Next, we sampled a larger set of models that are reasonable models of the VVS and asked: i) how well do these models perform when applied to amygdala responses and ii) what layer in each model is best predictive of amygdala responses (compared to which layer is the best predictor of IT responses). We first illustrate the examination of layer-wise prediction scores in Figure 2C, again using AlexNet, separately for IT and amygdala. Overall, response prediction performance increased with layer depth, similarly in both regions. We noted that the layer that best predicted the responses of amygdala sites tended to be from a slightly deeper layer than the layer that best predicted the IT sites (black arrows).

When we scaled this to test response predictions over thousands of layers across 78 task-optimized DNNs ([29]; TorchVision Models), we found that task-optimized models of the VVS can predict amygdala responses to natural images (Figure 2E green dots show best-performing layers for the models). Consistent with the direct anatomical connection from IT to the amygdala, we found that model layers that tended to be more capable of predicting the IT responses also tended to be more capable of predicting the amygdala responses (i.e. a strong correlation between the predictivity scores over all model layers; r = 0.98, p*<*0.001; see Figure 2E). However, closer examination revealed that, for amygdala response predictions, the most predictive model layer tended to be a slightly deeper layer than the most predictive IT layer (Figure 2D, layer depth is normalized between 0 and 1 for all models for comparability, median depth of maximal-amygdala-predicting layer was 0.75, c.f. 0.70 for maximal-IT-predicting layer, p *<* 0.001). Therefore, we conclude that with IT being our baseline, we can model the visual responses of some amygdala neurons to a reasonable degree, especially when using slightly deeper layers of models that perform well in IT (see also Discussion).

### Visuo-valent neurons in the primate amygdala

Next, given the amygdala’s multimodal inputs and role in emotional valence processing [5, 18, 33], we hypothesized that a subset of visually reliable amygdala units may also carry information about the valence of sensory events. To address this question, we assessed valence coding of amygdala neurons in 10 to 20 percent of experimental trials for monkey 1, by presenting a fixed tone-outcome sequence in which different tones indicated water reward, juice reward or airpuff as the outcome (see Methods).

Figure 3A shows responses of two example valence-coding cells that also respond to visual images. The top unit responds more strongly to airpuff than to juice and water reward. In contrast, the bottom unit responds more to juice and water reward than to airpuff. Notably, these units also reliably respond to visual input (Figure 3A, right), although these images were not explicitly associated with valence outcomes.

**Figure 3:**
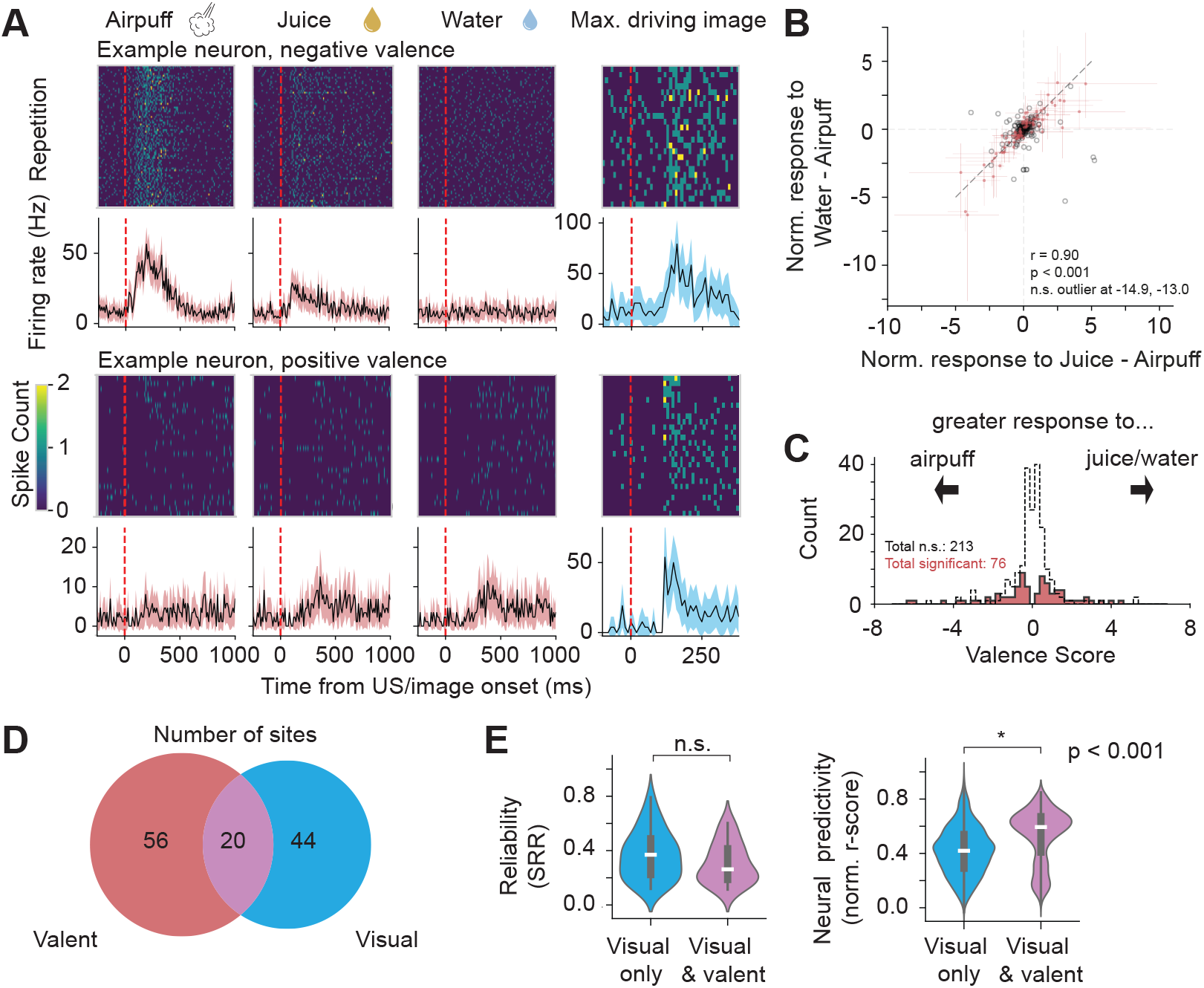
A) Example single neuron 1 that shows strong responses to airpuff and some visual images. Example neuron 2 that shows strong responses to juice and water, and some visual images. B) Valence neuron selection. Valence neurons are defined as those neurons where negative events (airpuff) can be reliably distinguished from positive events (water, juice, indicated in red, see Methods). For the majority of cells, water and juice responses contrasted in a similar direction to airpuff responses (most neurons are in the upper right and lower left quadrant). C) Distribution of valence scores for sites with (red) and without (black, open) reliable valence responses. D) Venn diagram of overlap between visual, valent and visuo-valent sites. E) Single-repetition reliability and neural predictivity of visual and visuo-valent neurons.

We found that a subset of amygdala units is “visuo-valent”, jointly encoding visual and valence information even in the absence of pairing through conditioning. A valence unit was defined as one that reliably differentiates juice vs. airpuff or water vs. airpuff (or both) (see Methods, [5, 19]). In valence units, normalized responses to juice vs. airpuff were highly correlated with normalized responses to water vs. airpuff (Figure 3B), supporting a common valence axis (r = 0.90, p *<* 0.001). Accordingly, a valence score was assigned to each unit based on the magnitude and direction of the normalized difference (Methods), with negative values for greater responses to airpuff (Figure 3C). Combined with our previous assessment of visual reliability, among 289 sites recorded with the full visuo-valent paradigm (i.e., including valence trials), 76 sites were classified as valent, 64 as visual, and 20 among these were both valent and visual (Figure 3D; 56/225 nonvisual sites and 20/64 visual sites were valent; *χ*^2^ test of independence: *p* = 0.39; 15 of the 20 visuo-valent sites were SUA).

Furthermore, the identified visuo-valent units were as visually reliable as visual-only neurons, despite being multimodal. Compared with sites that showed only reliable visual responses, those that additionally encoded valence had a comparable level of reliability (Figure 3E, top) and even higher ANN model-based predictivity (Figure 3E, bottom).

### Predicting valence preference cross-modally from visual tuning

So far, we established that some primate amygdala neurons show both high visual and valence selectivity. This raises the question of whether this association follows a systematic rule across the visuo-valent neural population, such that neurons encoding a specific direction of valence also prefer specific visual features.

We addressed this question by asking if the valence preference of each amygdala unit can be predicted from its visual preference using linear regression (Figure 4A, left; see Methods). To test for generalization among the population, linear regression models were constructed iteratively on all but one recording session and tested on held-out session data (Figure 4A, right). The predicted valence scores of held-out sessions correlated highly with the measured valence scores, with an *R*^2^ of 0.55 (p*<*0.001, Figure 4B). Note that this method is conservative, and leaving out sites instead of entire recording sessions results in an *R*^2^ of 0.82 (Figure S2). This finding implies that visual response preferences are linked to the valence preference of visuo-valent amygdala sites, in a way that is not idiosyncratic to each tested site. Notably, we find this even without ever explicitly pairing different images with different valence outcomes.

**Figure 4:**
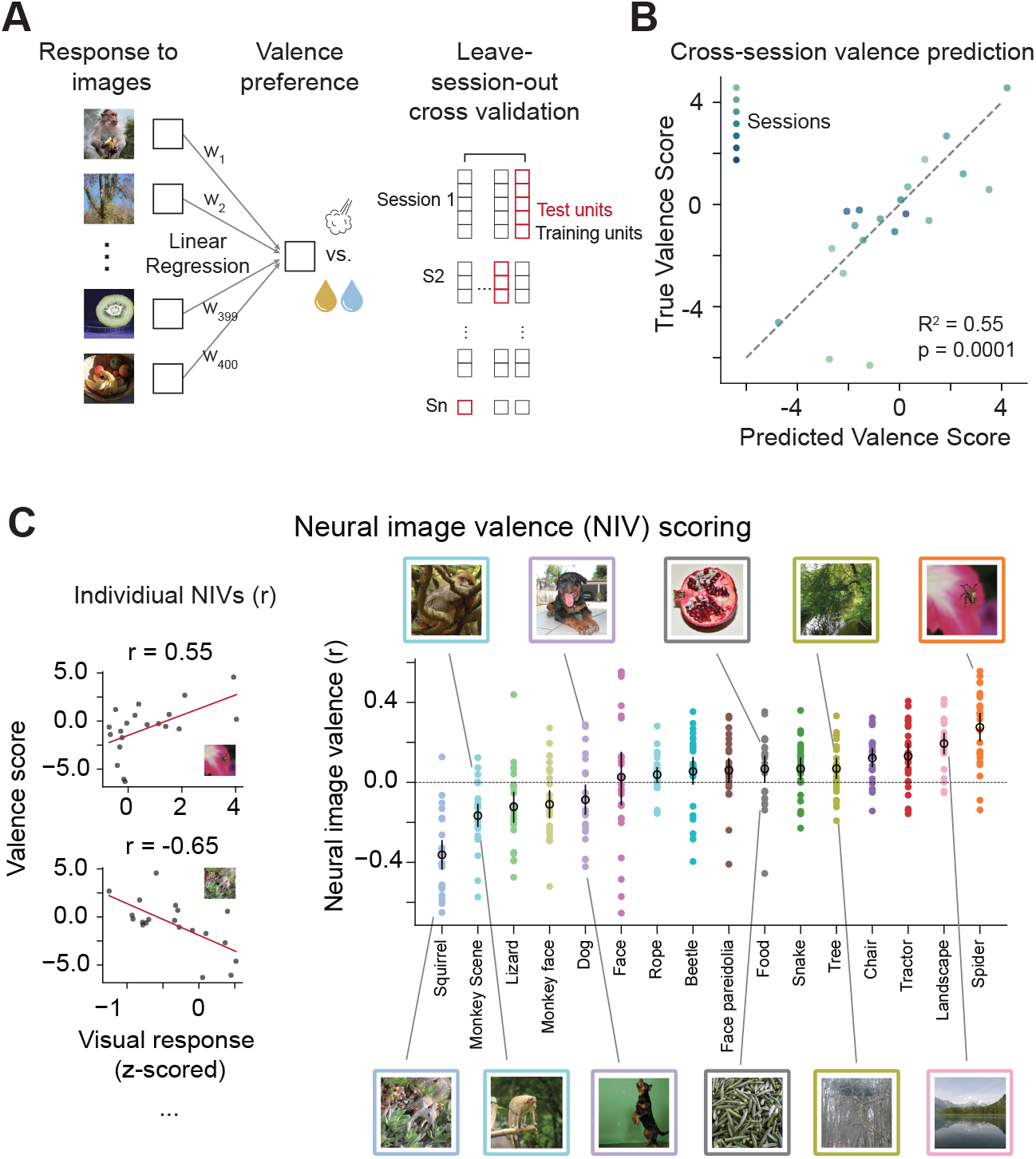
A) Illustration of regression modeling to predict valence based on visual selectivity. See also Methods. B) Actual versus predicted valence, model predictions of held-out recording sessions. Figure S2: Holding out recording sites instead of sessions improves performance. C) Left: Calculation of individual neural image valence (NIV) scores for example images. Right: NIV across categories and images, sorted by the average NIV per category. Black: Average NIV and SD per category. Positive values indicate images where a response is stronger for positive valence neurons.

Given the systematic visual-valence linkage found here, we examined the image content that tends to correlate with positive or negative valence responses. To this end, we defined each image’s neural image valence (NIV) as the correlation across sites between the image-evoked responses and the valence preference scores (Figure 4C, left, Methods). Here, an NIV value close to -1 indicates that the image elicits stronger activation for sites that respond more to airpuffs, whereas a value close to +1 indicates stronger activation for sites that respond more to liquid rewards. A one-way ANOVA revealed a significant category effect (*F* (16, 383) = 16.776, p*<*0.00001), with category explaining 41.2% of the variance in NIV, while substantial variability remained within categories, with both positive and negative NIV examples for many categories (i.e. above and below zero).

Within categories, NIV appeared to at least partially organize images in semantically meaningful ways. To note our visual impressions, negative human faces frequently show teeth and positive ones look away - which might make sense given that monkeys show teeth during aggression and fear, and eye contact is aggressive and known to correlate with airpuff responses in the amygdala. When images were post-hoc sorted by direct versus indirect gaze (as studied by [19]), images with indirect gaze had significantly more positive NIV (shuffle test, p*<*0.01). Also, positive-contributing images of mammals frequently show these in relaxed posture and/or eating, fruit images tend to be greener when more negative, trees less leafy. Although a rigorous exploration and validation of these patterns lie beyond the scope of this paper, we provide the NIV-sorted images per category in Figure S3 for the interested reader.

We conclude that valence preference of visually-driven amygdala neural sites can be at least partially predicted from their image-driven responses alone, and that there is some lawful connection between preferred visual and non-visual inputs even when all images are repeatedly paired with an identical outcome (other images and water reward outcome, see also Discussion).

### ANN models yield predictions of valence for any image

Finally, we asked if this association between visual image content and valence preference could be captured using ANN models of the VVS, given our previous finding that good ANN models of IT responses also have predictive power for visually-driven amygdala responses.

We found that the NIV of each image could be directly predicted from its pixels using ANN models of the VVS (Figure 5A), once the relationship between image and valence was established using cross-validated ridge regression (see Methods). Indeed, the per-image NIV could be predicted by deep layers of ANNs (Figure 5B), especially for models that performed well in predicting IT responses as well (Figure S4). This suggests that the visual representations underlying visual-valence association in the amygdala share a common basis with those in IT.

**Figure 5:**
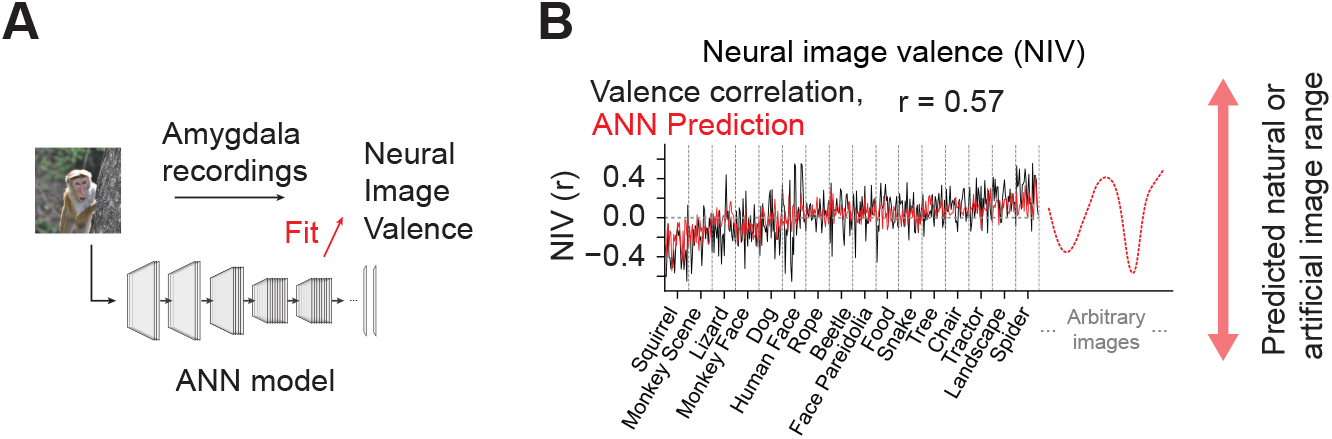
A) Image valence prediction based on example deep neural network model by layer. B) Neural image valence (NIV) measured (black) versus artificial neural network prediction (red). Toward the right, illustration of possible uses of the model.

Notably, because the model operates on pixels, the model can make NIV predictions on arbitrary images, i.e. including natural images not tested, different categories, and artificial stimuli, based on the observed structure of the data and the learned image statistics of the model. Such models and their predictions could be used in a number of ways, by identifying promising new stimuli to try, by bridging between subjects and experiments using different stimulus sets, and by generating artificial stimuli that are predicted to drive valence responses in an intended direction.

## Discussion

In summary, we provide evidence that the primate amygdala has strong, selective visual drive, and that it non-uniformly links that visual drive to valence encoding, even without explicit conditioning. Moreover, that visual drive and associated valence coding can be reasonably well predicted by simple extensions of contemporary, machine-computable models of the ventral visual stream (VVS). Specifically, electrophysiological recordings revealed that some amygdala sites responded reliably and selectively to brief (*∼* 100 *ms*) image presentations of a broad range of different image categories, with reliable selectivity comparable to IT sites. That is, to first pass, the visual drive of many amygdala neurons is indistinguishable from that of IT neurons – many of which provide direct axonal input to the basolateral amygdala. Note that our recordings were targeting and therefore biased toward the lateral aspects of the basolateral amygdala, but that we refer to amygdala sites since sampling from other parts of the amygdala cannot be excluded. Also similar to IT, these image-driven amygdala responses were quantitatively predictable to reasonable accuracy (predictivity *r ∼* 0.5 for the best-performing model-layer combinations) using extensions of contemporary ANN models of the VVS, with amygdala sites best captured by slightly deeper model layers than those optimal for IT.

A subset of visually reliable amygdala sites also encoded valence, with some neurons preferring unconditioned positive events (US+) and others preferring unconditioned negative events (US-). This unconditioned valence coding is presumably carried via other anatomical pathways into the amygdala [18, 34, 35], but we found that it was not unrelated to the aforementioned visual selectivity. On the contrary, for this subset of sites, specific amygdala neural visual preferences were correlated with preferences for positive versus negative valence events. That is, if one only knows the visual selectivity pattern of one of these neurons, one can reasonably accurately predict the valence preference of that neuron (*R*^2^ *∼* 0.55, Figure 4B, conservatively estimated). This suggests that the pattern of visual drive carried by the connections from IT to positive-preferring and negative-preferring amygdala sites is not uniform or random, but that it is instead biased, naturally linking some types of visual drive to US+ neurons and other types of visual drive to US-neurons, without any explicit differential temporal-association conditioning of visual stimuli.

Finally, ANN-based models predicted which images cause strong responses in positive or negative valence neurons, enabling quantitative prediction of neural valence states in the primate amygdala directly from pixels.

Where does this visual-valence linkage come from? The visual-valence linkage observed here is unlikely to be a learned artifact from our laboratory experiment. In our task, all images in the RSVP stream were equally predictive of the same water reward at the end of a trial, making it implausible that differential image responses reflect differences in experimental reward expectation (in contrast to e.g., [13, 15]). This equal predictivity was established through several weeks of training sessions before recording, which also means that all images were equally familiar [15, 36]. The rapid and randomized presentation further reduced potential confounds arising from free-viewing, stimulus anticipation, or changes in attention [37, 38]. Thus, the observed visual–valence linkage in the amygdala likely reflects a pre-existing relationship — either innate, shaped by lifelong visual experience, or a combination of both — rather than one induced by laboratory training. This notion is consistent with computational studies showing that high-level functional tuning can emerge in ANNs from architectural constraints of the VVS and optimization on ecologically relevant tasks, without explicit training on the target attribute [39–42].

Our results support the view that the amygdala can be considered a generic visual processor with lasting biases, alongside its well-known role as a learning hub linking stimuli to outcomes. First, the top-driving images of visually reliable neurons belonged to a broad range of categories, suggesting that the amygdala encodes rich visual features (Figure 1). This is consistent with two of the most comprehensive studies of visual responses in the non-human primate amygdala, based on both fMRI and single-unit recordings, as well as studies in epilepsy patients which report complex responses across diverse image categories [10, 11, 43–45] beyond traditionally emphasized stimulus classes such as faces [12].

Second, ANN models trained for core object recognition captured substantial variance in amygdala visual responses, reaching predictivity up to *r ∼* 0.5 for the best-performing layers (Figure 2E). This indicates that, despite the fact that the amygdala is anatomically not a purely neocortical structure and is best known for learning stimulus-outcome associations, it preserves a robust visual processing structure that is well approximated by VVS computations (see also recent findings [44, 46]). Third, the best-predicting ANN layers for the amygdala tended to be slightly deeper than those for IT, positioning the amygdala as a downstream stage closely following IT. This is also in line with the suggestion that MTL structures form the pinnacle of the visual hierarchy as suggested by [47]. Together, these findings help reconcile a divide between “visual feature-based” and “conceptual/semantic” accounts of amygdala responses in humans [44, 45], by suggesting that they reflect a continuum rather than mutually exclusive functions.

“Valence” was operationalized along a single outcome-based dimension in this study, while richer representations may emerge with a broader set of affective events. In our analysis, we summarized each site’s valence preference with a single scalar “valence score”, defined as the site’s normalized response difference between negative (airpuff) and positive (juice or water) outcomes. This 1-D definition of valence is supported by the strong correlation between the juice-airpuff and water-airpuff differential responses and is consistent with prior work defining valence from such US responses [5, 19]. However, we do not preclude the possibility that affective outcomes are represented along multiple dissociable axes depending on event type (e.g., different classes of threat such as predator cues, or different reward types such as social reward) [3, 16]. Accordingly, the valence neurons here are likely a subset identified under a limited set of valence events, and sampling a broader range of outcomes could reveal additional valence-coding neurons, which may or may not also have a non-conditioned relationship to visual coding.

Similarly, “neural image valence” NIV was operationalized as a systematic correlation between valence and response strength to an image across the neural population. This allowed us to determine which image content is most responsible for the observed visual-valence predictivity. This constitutes a novel way to assess the valence of images, by determining the biases in the association between image and valence responses across a large set of images. Typically, image valence is assigned by human consensus report (e.g., OASIS, IAPS databases [48, 49]), estimates of human experts when it comes to other primates [11, 50], or is imposed through conditioning. Potential links between NIV and the gold standard of human report remain to be established. However, it is clear that the NIV metric is based directly on responses of (and thereby inherently relevant for) the amygdala, it scales, and is assessed cross-modally, linking images to other events. We therefore consider this metric a promising avenue for affective science, especially when combined with image-computable models.

Image-computable models can provide a shared quantitative framework for affective science, bringing several key advantages. One advantage is the direct comparison of different alternative models, and a comparison of modeling success with that of other brain areas. This is particularly valuable given our observation that models that better explain VVS patterns of visual drive also tend to better predict amygdala patterns of visual drive, so progress in one domain can directly benefit the other. Next, because only a limited set of stimuli can be tested in any single experiment, models provide a principled way to generalize beyond the specific images used, addressing long-standing concerns about what conclusions extend beyond selected stimulus sets [10, 11]. In practice, models can generate quantitative affective predictions for arbitrary, untested images or categories, allowing researchers to formalize and compare competing hypotheses. Moreover, the models themselves constitute a shareable, explicit hypothesis that can be tested against new data across images and individuals, facilitating cumulative validation and refinement across experiments.

Finally, our results suggest a path toward non-invasive modulation of limbic circuits. Because our models operate directly on pixels, they can predict how arbitrary images are expected to drive positive- or negative-valence neurons in the primate amygdala. This enables the synthesis of visual stimuli designed to optimize the activity of neuronal subpopulations associated with e.g. a specific valence direction. Indeed, prior work has shown that ANN-based models of the VVS can be not only used to predict, but also to systematically control image-driven neural activity by synthesizing images that selectively drive targeted neural populations [22, 23, 51]. Extending these approaches to the amygdala, our results suggest a potential route for modulating valence-related neural activity through controlled visual input. Given that aberrant amygdala neural activity is implicated in a range of disorders [52, 53], such a non-invasive, model-guided strategy could provide a promising direction for potential future therapeutic intervention.

## Acknowledgments

We thank T. Pistone, S. Goulding, and T. Fakhoury for assistance in animal care and training as well as surgical assistance and Y. Bai and K. Kar for contributions to surgical planning and execution and the DiCarlo lab for helpful discussions.

## Supplemental Figures

**Figure S1:**
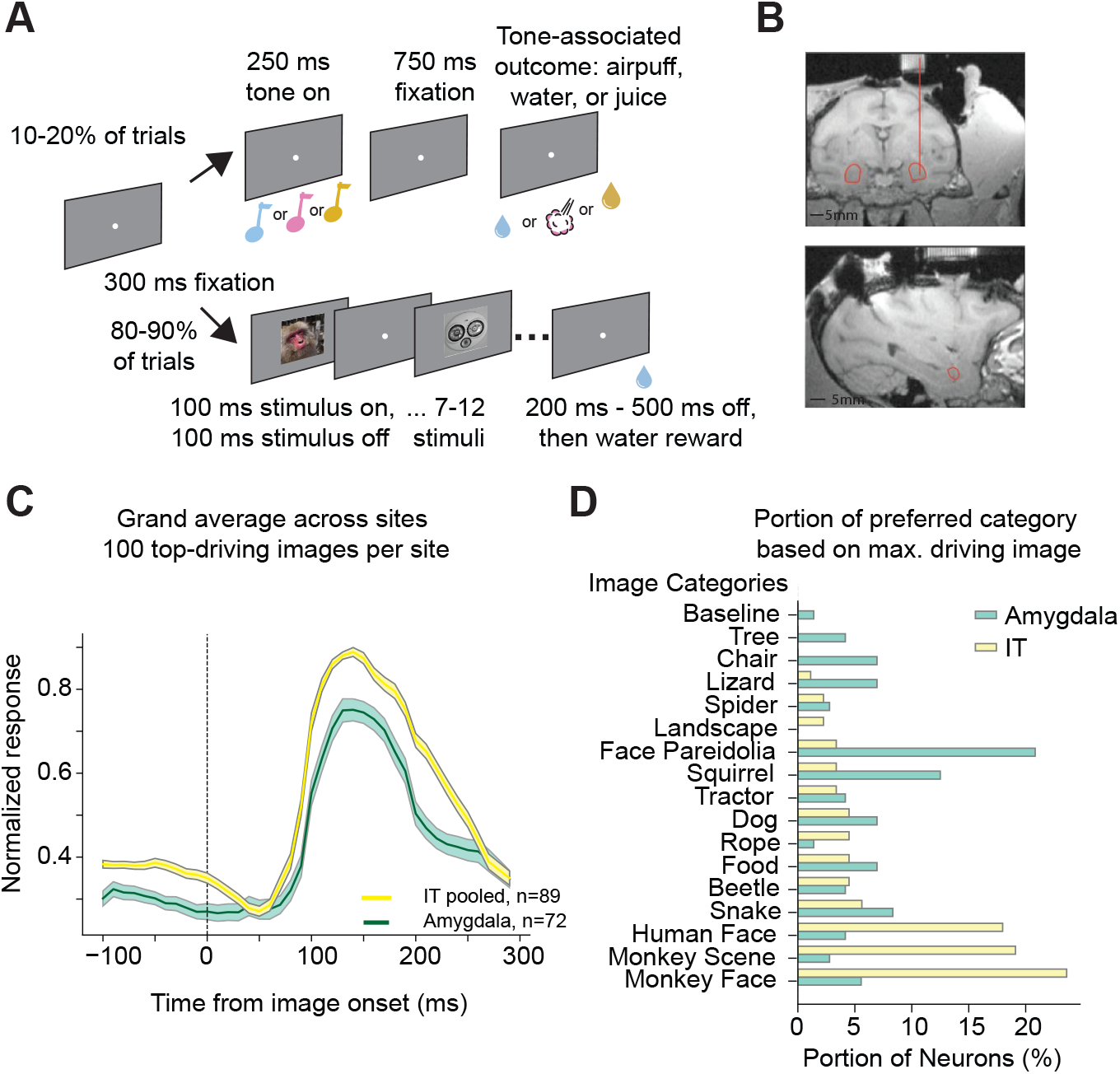
A) Schematic of the task for monkey A with amygdala recordings. B) Structural MRI scan of monkey A together with recording grid across coronal (top) and sagittal (bottom) anatomical planes. Recordings targeted lateral parts of the amygdala (lateral and basal nuclei, AP coordinates 20, 21). Red vertical line: projection from recording grid hole to target region. Red outline: amygdala based on [54] C) Grand average, normalized response to image onset. For each site, responses to the top-10 driving images were selected. Responses were averaged across images, then normalized to the peak and then averaged across sites. SEM is shown across sites. See also Methods. D) Portion of neurons responding with their maximal response to a particular image of each category.

**Figure S2:**
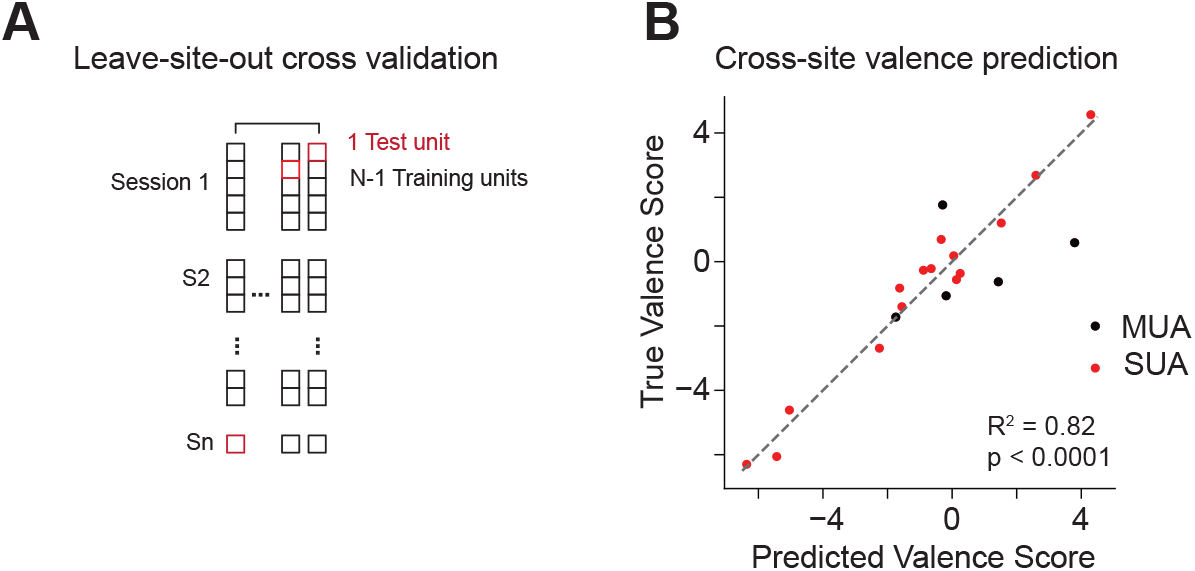
A) Leave-one-site out based prediction of valence based on visual response preferences, see Figure 4B for leave-one-session-out predictions. Dark dots indicate MUA sites, brighter dots SUA sites.

**Figure S3:**
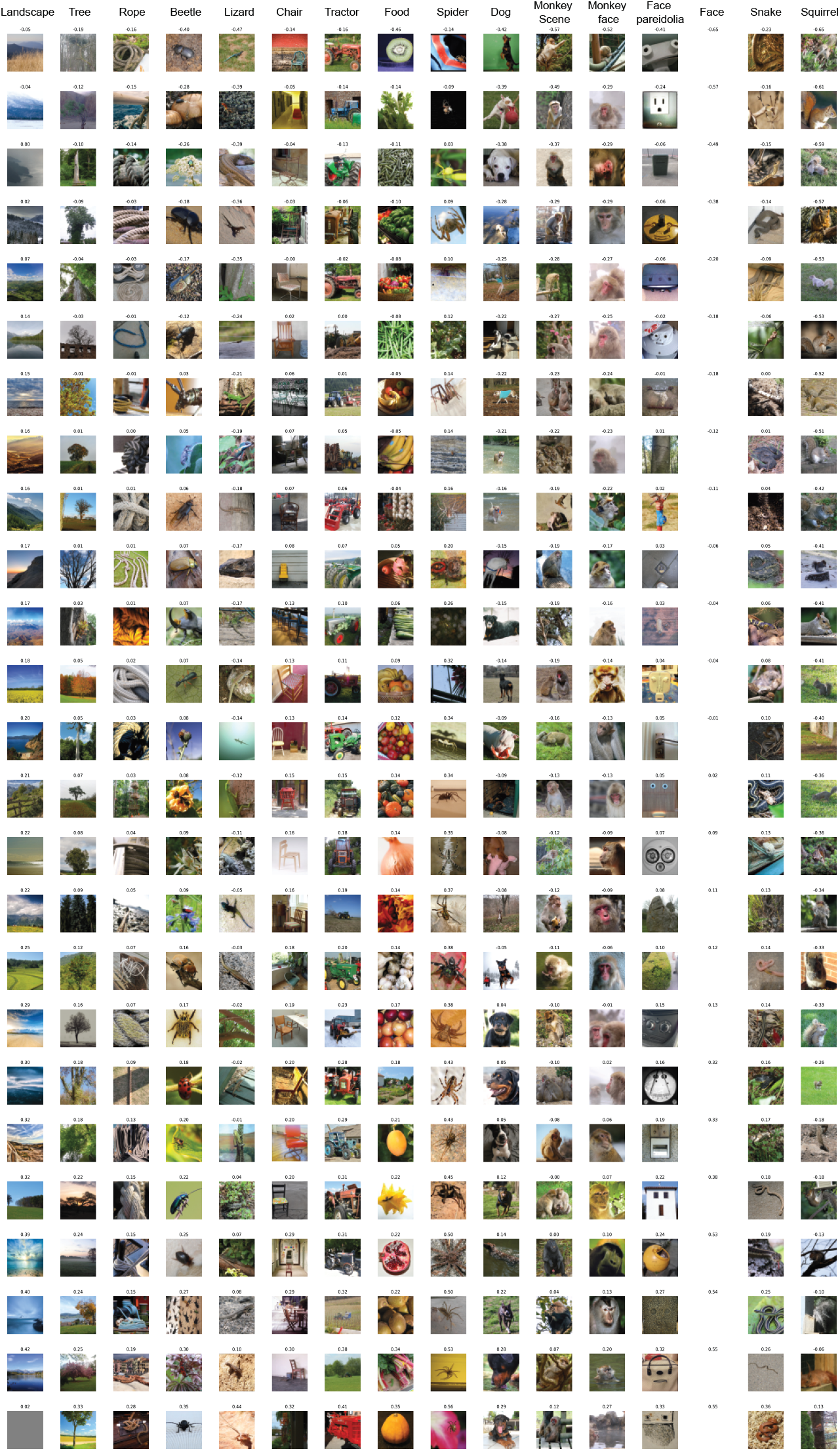
Individual images per category and their respective NIV score. Sorted within category by NIV from most negative (top) to most positive (bottom). Images of persons have been removed for the purposes of publication only and will be made available on a secure database, available on request.

**Figure S4:**
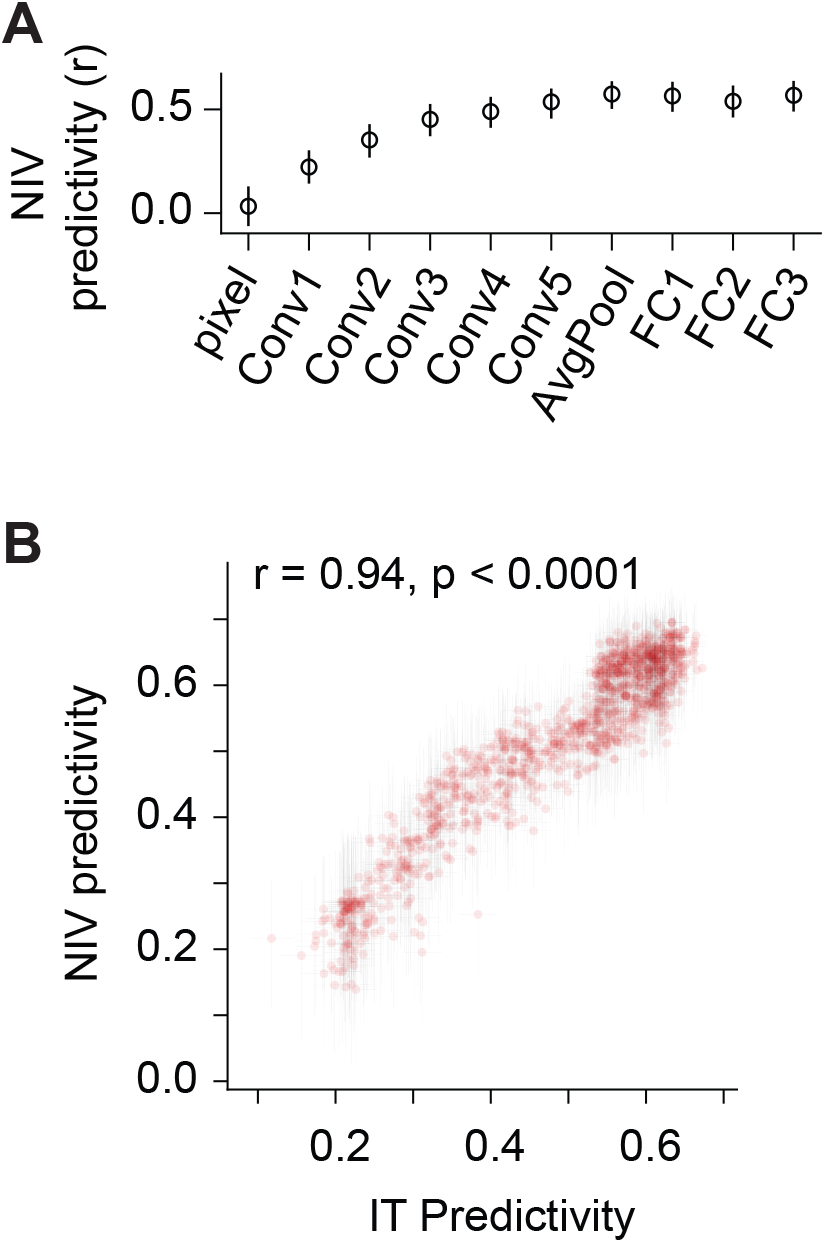
A) Predicted NIV across layers for an example ANN (AlexNet). B) Correlation of IT predictivity with NIV predictivity across models.

## Methods

### Task

For the IT animals, the task consisted only of visual trials. For the amygdala monkey, the task consisted of randomly interleaved trials of two different basic kinds, which we will call visual trials and valence trials. In both cases, a white fixation point (0.2 dva) would light up and the animal self-initiated a trial by fixating within a 2 dva window for 300 ms. Visual trials: A series of rapidly presented visual stimuli were shown 300 ms after fixation onset (RSVP). Individual images (described in section Images) were presented for 100 ms with 100 ms blank gray screen in between, for 7–12 images per trial (sampled uniformly). This was followed by another blank period of 200–500 ms and reward delivery (water) if completed without a break in fixation. A break of at least 500 ms was enforced before the next initiation of a trial was possible. The different images occurred in random order, and repeated in a new random order once each image in the set was shown, for a median of 20/27/45 repetitions (M1/M2/M3) repetitions per image and session. RSVP trials follow the same design as typically used for our IT studies [31, 55, 56]. This design allows us to acquire many images and repetitions for model fitting. Valence trials: In the amygdala animal, 10–20% (depending on session) of trials (selected at random) were valence trials. In valence trials, one of three sinus tones would play for 250 ms, 300 ms after fixation onset (baseline period in analyses). There was a further 750 ms fixation period, followed by a deterministic outcome. These outcomes, depending on the tone, were either a water reward, a juice reward of equal duration and magnitude, or delivery of an airpuff (10 PSI, 150 ms) to the face area. Trials repeated on error, since the animal had a tendency to break off trials with an airpuff tone. The animal was highly familiar with the tone-outcome mapping at the time of recordings. We call these valence trials based on previous naming of “valence cells” and agreement in the literature that airpuffs of sufficient strength are negative, whereas water and juice are mildly positive and positive outcomes respectively [5, 19]. They were intended to “ping” the nature of the cells recorded. Tasks were delivered with mWorks (v12) and presented on an LG Ultragear 32GP850-B monitor (2560 x 1440 pixels) at 120 Hz and 50 cm distance to the animal.

### Images

Natural images were collected from open databases on the internet (Wikimedia Commons, Yahoo Flickr Creative Commons 100 Million (YFCC100M) [57], Kaggle). About tenfold more images per category were collected than used during the recordings, and the sample images for the final set of 400 images (25 per category) were drawn at random from these larger collections. The categories spanned inanimate (tractor, rope, chair, face pareidolia), non-primate mammals (dog, squirrel), and other animals (snake, lizard, spider, beetle), food (fruit and vegetables), nature scenes (trees and landscapes) and monkey (macaque) scenes, as well as monkey and human faces. These categories span those that have been deemed important to amygdala or valence responses by previous research (faces, conspecifics, face pareidolia, snakes, spiders) [12, 50, 58] as well as stimuli that share some visual (and/or semantic) properties with these categories (squirrels, dogs for monkeys, lizards, ropes for snakes), as well as intentionally arbitrary categories such as tractors and chairs. In addition, the screen was shown blank in background color (baseline gray) 1/401 times at random, as frequently and in the same manner as other images. Images of human faces were removed for the purposes of publication only.

### Surgical implants and electrophysiological recordings

All procedures comply with the NIH guidelines for the use of non-human primates in research and have been approved by MIT’s Institutional Animal Care and Use Committee. All monkeys were surgically implanted with a head-post under aseptic conditions. One adult male rhesus macaque, monkey 1 (weight 12 kg; age 7 years), was prepared for neurophysiological recordings from the amygdala. Two adult male monkeys (monkey 2/3, age 10/10 years, weight 11/10 kg) were prepared for IT recordings, both from the left hemisphere.

The stereotaxic coordinates of the right and left amygdala were determined based on high-resolution 3T structural magnetic resonance imaging (MRI) scans (voxel size = 0.5 mm isotropic). For each hemisphere, a craniotomy was made and a rectangular (21 × 24 mm inner dimensions) polyether ether ketone (PEEK), MRI compatible recording chamber was surgically attached to the skull (the first chamber was explanted when the second chamber was implanted).

In IT, neural activity was recorded with 10×10 microelectrode arrays (Utah arrays, Blackrock Microsystems), as previously described [56]. Each array contained 96 electrodes (excluding corner electrodes), each 1.5 mm long and with a 400 µm spacing between electrodes. The placement of arrays was guided by the visible sulcus pattern during surgery. Arrays sampled a variety of regions along the posterior-to-anterior IT axis. Nevertheless, for all analyses, we refrained from taking the specific spatial location of the electrodes into account, treating each site as a random sample from the overall IT population. One session was recorded per animal.

All recordings were made with a 512ch Intan recording controller and RHX data acquisition software v 3.1.0, 30 kHz sampling rate, 0.79 Hz to 7.6 kHz hardware bandwidth. In the amygdala, neural activity was recorded with linear electrode arrays (V-probes, 100 micron spacing, 16/24 channels, Plexon Inc, Dallas, TX [7 sessions] and DBC-128-2 Deep arrays, Diagnostic Biochips, 50 micron spacing for two 64 channel arrays 40 micron apart, shifted so that contacts are 25 micron distance vertically to their lateral neighbor [8 sessions]). A single electrode array was acutely lowered into the amygdala for each recording session using a Narishige hydraulic drive (Narishige, Tokyo, Japan). During recordings, a grid with 1 mm distance between cannula guide holes was placed in the chamber, a twenty-three-gauge cannula was inserted through the guide holes and placed approximately 6 mm into the cortex. Probes were manually advanced to approximately the same depth as the guide tube and then down to the amygdala at a rate of 15–20 *µ*m/s, slowing to 5 *µ*m/s after the tip of the probe was within a few millimeters of the start of the amygdala. Data from a total of 15 sessions were analyzed.

### Spike sorting

Spike sorting was performed using Kilosort 2.0 and default settings (150 Hz high pass filter, min. firing rate 1/50 Hz in a cluster, amplitude penalty 10, AUC for cluster split 0.9, projection thresholds [10 4], spike threshold -6 std, drift correction), followed by manual inspection for the following criteria: amplitude above the noise consistently for the recording, isolation distance to nearby clusters, clean refractory period by Kilosort and phy (https://phy.readthedocs.io) evaluation [59]. 351 units were retained for analysis, 215 of them multiunits. Data analysis is pooled across single- and multiunits unless otherwise noted. In IT, all detected units were multiunits (M2: 141, M3: 64). Therefore, we will refer to multi- and single-units as “neural sites”, and pool single- and multiunit activity in the amygdala unless otherwise noted.

### Eye tracking

During recording sessions, we monitored the monkeys’ eye movements using video-based eye tracking. At the start of a session, monkeys performed an eye-tracking calibration task by making a saccade to spatial targets and maintaining fixation for 500–700 ms. A single eye was recorded with Eyelink1000 Plus v6.09 at 500 Hz.

#### Identifying visually reliable neural sites

#### Single Repetition Reliability (SRR)

For each neural site, the per-trial spike counts to visual stimuli were averaged over a 100– 200 ms analysis time window for the amygdala and a 70–170 ms for IT, and then converted to firing rate (spikes/s). The window length was chosen based on the duration of the stimulus (100 ms), and is typical for IT recordings, and the start time was determined based on the earliest reported visual response latencies for primate amygdala and IT neurons [approximately 100 ms and 70 ms, respectively; amygdala: [10], IT: [60]].

The SRR for each neural site was calculated by sampling the time-averaged responses from two random repetitions per image and computing the Pearson correlation between the first and second responses across all images. This process was repeated 1,000 times and the average correlation was taken as the SRR of this site.

Neural sites were identified as visually reliable if the lower bound of the 95% confidence interval of the sampled SRR distribution exceeded zero.

For reliable sites, a control analysis was performed. SRR was computed with one of the repeats drawn randomly from the first half of the recording session and the second repeat randomly from the second half, instead of selecting 2 repeats fully at random, to assess stability over the recording session. Note that due to the pseudorandom presentation, this means that several thousand images have been shown between these repeats on average.

### Split-Half Reliability (SHR)

The SHR for each neural site was calculated by randomly splitting the repetitions of each image into two halves, averaging the responses within each half, and calculating the Pearson correlation between the two response vectors across all images. This process was repeated 1,000 times, and the average correlation, after applying the Spearman-Brown correction, was taken as the SHR of this neural site.

### Identifying valence neural sites

#### Valence score

For each neural site, spike responses were measured across trials for juice, water, and airpuff events, with *n*_*juice*_, *n*_*water*_, and *n*_*airpuff*_ repetitions, respectively. Per-trial spike counts were averaged over the 100–500 ms response time window following each event and normalized by the baseline activity during the pre-CS presentation period (-250–0 ms) as follows:

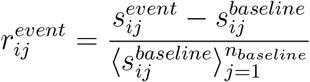

where 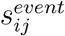 represents the average spike count of site *i* on the *j*-th repetition of a given event type, and 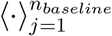 denotes averaging across repetitions *j* = 1 to *n*_*baseline*_.

The valence score for each neural site was defined as the difference in the average normalized response between one of the reward events (juice and water) and the aversive event (airpuff) as follows:

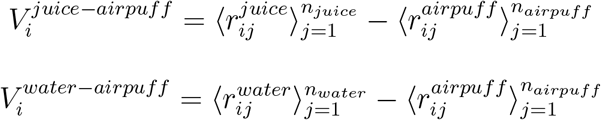

### Reliability of valence encoding

To estimate the reliability of the valence encoding for each neural site, bootstrap samples of the neural site’s spike responses were generated by randomly sampling trials (with replacement) for each event, using the same number of trials as the original repetitions. The valence score was recalculated for each bootstrap sample. This process was repeated 1,000 times, resulting in a distribution of 1,000 valence scores for each comparison.

Neural sites were identified as valence sites if the 95% bootstrapped confidence interval of the valence score excluded zero (i.e., the lower bound is greater than zero or the upper bound is less than zero). The confidence interval analysis was applied to both juice vs. airpuff and water vs. airpuff comparisons, and the union of sites significant in either comparison defined the final valence set. If a neural site showed significant valence scores in both comparisons, the score with the greater absolute value was taken as its valence score. Each valence neural site was classified as positive or negative according to the sign of its valence score.

### Visuo-valent neural sites

Neural sites identified as both visually reliable and valence-encoding were defined as visuo-valent neurons.

## Analysis of visual encoding

### Average visual response profile and spike counts

For each neural site, the average visual response profile was obtained by averaging the visual PSTH within the response time window (70–170 ms for IT and 100–200 ms for the amygdala) and across repetitions. This resulted in a 400-dimensional response vector (one value per image) for each neuron. Time-resolved data with spike counts is shown in 10 ms bins (e.g. Figure 1C). The findings for Figures 1 and 2 held also if the IT analysis window was set to the same time range as the amygdala time window.

For Figure S1C, the grand average responses per recording region, the grand average, normalized response to image onset was computed. For each site, responses to the top-10 driving images were selected, based on a shared 100–200ms time window. Responses were averaged across images, then normalized to the peak. IT sites were pooled and then averaged across sites. SEM is shown across sites. Conclusions remained similar when changing the number of top-driving images or using the 70–170ms time window.

### Predicting visual responses using task-optimized ANNs

#### a. Regression model

Building on previous work modeling the ventral visual stream [21, 31, 40], we tested whether visual features (**z**) extracted from ANNs trained for core object recognition could predict each neural site’s visual responses to images as follows:

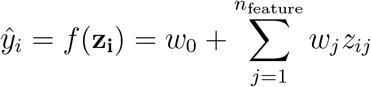

where 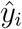 is the predicted response of this neural site to image *i, w*_0_ is the intercept, *w*_*j*_ is the coefficient for the *j*th feature, and *z*_*ij*_ is the *j*th feature value of image *i*.

The model was constructed using ridge regression that minimizes the following loss function:

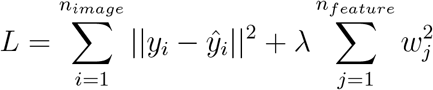

where *y*_*i*_ and 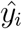 are the observed and predicted responses to image *i, λ* is the regularization parameter, *n*_*image*_ is the number of images used for model training, and *n*_*feature*_ is the number of features extracted from the image.

#### b. Feature extraction

For image feature extraction, we used 78 pretrained ImageNet-1K classification models from the PyTorch framework, spanning CNN and Vision Transformer families (e.g., AlexNet, VGG, ResNet, ViT). Features were extracted from all major layers of each architecture (i.e., the top-level stages in the Torchvision module hierarchy), yielding 1,235 layer-wise feature sets in total.

Given that our dataset consists of 400 images, the number of features in the intermediate layers of a typical ANN can be excessively large. To address this, we used the sparse random projection method (python scikit-learn package) to map the activations from each layer of pretrained DNN models to a lower-dimensional space. This method compresses high-dimensional data into a low-dimensional space with a fixed dimensionality (*n*_*dim*_ = 5,135), determined by the Johnson-Lindenstrauss lemma, while preserving pairwise distances up to a small error margin (*ϵ* = 0.1).

#### c. Training and testing

Using the feature extraction and regression framework described above (a-b), a ridge regression model was trained for each amygdala/IT neural site to predict its responses from the projected image features of each model layer. Nested cross-validation was employed to prevent overfitting. In the outer loop, the 400 images were split into a training set of 320 images and a test set of 80 images (5-folds). Leave-one-out cross-validation (scikit-learn RidgeCV function) was used on the training set to find the optimal ridge regularization hyperparameter (*λ*). The model with the optimal hyperparameter was trained on the training dataset and used to predict the responses on the test set. The prediction was repeated for each outer loop, resulting in response predictions for all 400 images. Model predictivity scores were calculated as the Pearson correlation between the predicted and true responses for each neural site. To account for measurement noise, a normalized model predictivity score was computed by dividing this correlation by the SHR of the same neural site, which serves as the theoretical upper bound on achievable predictivity. The best-performing layer of a model was defined as the layer showing the highest mean normalized predictivity across neural sites.

## Association between visual and valence coding

### Predicting valence score from visual response profiles

To investigate the potential association between visual and valence tuning in the amygdala, we tested whether the valence preference of visuo-valent neural sites can be predicted from their visual response profiles using principal component analysis (PCA) followed by a linear regression.

#### a. Data preprocessing and dimensionality reduction

Let *X ∈ R*^*n×m*^ be the response matrix (*n* = 72 visually reliable neural sites, *m* = 400 images). For each neuron *i*, the responses were standardized by subtracting the mean (*µ*_*i*_) and dividing by the standard deviation (*σ*_*i*_) across the images.

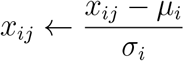

Next, we applied PCA on *X* for dimensionality reduction. The top *k* = 20 principal axes (explaining *≈* 75% of the variance) defined the image loadings *L*_*k*_ *∈ R*^*m×k*^. Embeddings (*Z ∈ R*^*n×k*^) were obtained as

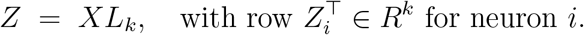

To avoid overfitting, the number of principal axes was matched to the number of visuo-valent neurons used in the subsequent regression (*n*_val_ = 20; thus *k* = 20).

#### b. Regression

Let 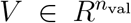 be the observed valence scores for the visuo-valent subset, and let 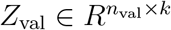 be their embeddings. We modeled each site’s valence score as

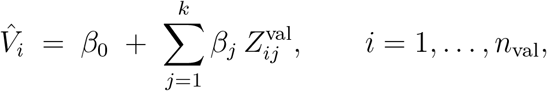

where *β*_0_ is the intercept, *β*_*j*_ is the coefficient for the *j*-th principal component, and 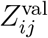 is the *j*-th PC coordinate of neuron *i*.

The model is constructed using ridge regression that minimizes the following loss function:

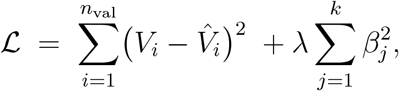

where 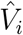 is the predicted valence score of the site, and *λ* is the regularization parameter determined through cross-validation (see below).

#### c. Nested cross-validation

To ensure robust model evaluation and hyperparameter selection, we employed nested cross-validation with two alternative outer-loop schemes:

Leave-one-site-out: In each outer iteration, data from one site were held out for testing, and data from the remaining sites were used for training. Within the inner loop, leave-one-out cross-validation was conducted on the training data to find the optimal *λ* (from 10^*−*5^ up to 10^4^, powers of ten). Using this *λ*, the ridge model was refit on the full training set and used to predict the valence score of the held-out site. This process was repeated so that every site served once as the test site, yielding valence-score predictions for all sites.

Leave-one-session-out: To further test if this association was shared across spatially non-localized neurons, in each outer iteration we held out all sites from one recording session as the test set and trained on sites from the remaining sessions. As before, *λ* was selected by leave-one-out cross-validation within the training sites, after which the model was refit on the full training set and used to predict the valence scores of the held-out session’s sites. This was repeated across sessions so that every session served once as the test fold.

#### d. Evaluation of significance

After completing the nested cross-validation, we concatenated all outer-fold valence score predictions 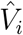 and their held-out targets *V*_*i*_ and computed the coefficient of determination.

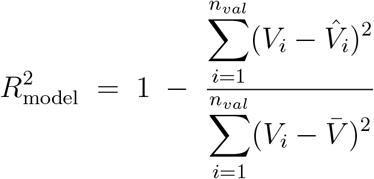

Here, 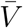 is the mean of all target valence scores.

We compared the model’s performance with a baseline that, in each outer fold, predicts the valence of all test neurons as the mean valence of the training neurons in that fold. This baseline ignores visual responses and serves as a trivial comparator. For each valence score prediction of neuron *i*, we computed the per-neuron squared error (SE)

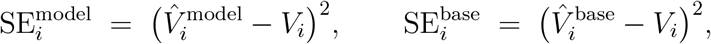

and defined the performance difference

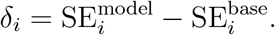

We then obtained a bootstrap 95% confidence interval for the mean performance difference 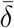 by resampling *δ*_*i*_ with replacement and concluded significance when the CI excluded 0 (upper bound *<* 0).

### Neural image valence

Let 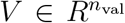 denote the valence scores of the visuo-valent amygdala neurons, and let 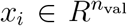 be the standardized firing activities (*X*) evoked by image *i* across the same neurons. We define the neurally defined valence of image *i* as the Pearson correlation between these two vectors:

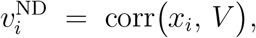

Values of 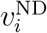 close to +1 indicate that neurons with more positive valence scores tend to respond more strongly to image *i* (and those with negative valence respond more weakly), whereas values close to *−*1 indicate the opposite pattern. Values near 0 indicate no systematic association.

### Predicting neural image valence using task-optimized ANNs

Identical to the visual response prediction using ANNs, we modeled each image’s neurally defined valence from ANN layer features,

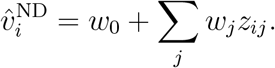

Training/validation and performance evaluation followed the identical protocol described above.

